# Quantifying the relationship between pollinator behavior and plant reproductive isolation

**DOI:** 10.1101/094896

**Authors:** Robin Hopkins

## Abstract

Pollinator behavior is an important contributor to reproductive isolation in plants. Despite hundreds of years of empirical research, we lack a quantitative framework for evaluating how variation in pollinator behavior causes variation in reproductive isolation in plants. Here I present a model describing how two aspects of pollinator behavior – constancy and preference – lead to reproductive isolation in plants. This model is motivated by two empirical observations: most co-occurring plants vary in frequency over space and time and most plants have multiple pollinators that differ in behavior. These two observations suggest a need to understand how plant frequency and pollinator frequency influence reproductive isolation between co-occurring plants. My model predicts how the proportion of heterospecific matings varies over plant frequencies given pollinator preference and constancy. I find that the shape of this relationship is dependent on the strength of pollinator behavior. Additionally, my model incorporates multiple pollinators with different behaviors to predict the proportion of heterospecific matings across pollinator frequencies. I find that when two pollinators display different strength constancy the total proportion of heterospecific matings is simply the average proportion of heterospecific matings predicted for each pollinator. When pollinators vary in their preference the pollinator with the stronger preference disproportionally contributes to the predicted proportion of total heterospecific matings. I apply this model to examples of pollinator-mediated reproductive isolation in *Phlox* and in *Mimulus* to predict relationships between plant and pollinator frequency and reproductive isolation in natural systems.

## INTRODUCTION

The notion that pollinator behavior is important for plant diversification and speciation can be traced to Darwin’s foundational ideas of evolution by natural selection (Darwin 1862). For many angiosperms, animals are necessary vectors for pollen movement between outcrossing individuals (Ollerton et al. 2011). It has long been argued that plants have evolved elaborate floral traits, including variable size, color, shape, and scent to attract and influence the behavior of pollinators (Campbell et al. 1997; Galen 1989; Schemske and Bradshaw 1999; Stanton et al. 1986). Furthermore, there is phylogenetic evidence that diversification in angiosperms is driven, in part, by pollinators (Valente et al. 2012; Van der Niet and Johnson 2012; Whittall and Hodges 2007). From a plant’s perspective, pollinator visits are only fruitful if the pollinator moves pollen between conspecific plants. Heterospecific pollen movement can result in hybridization, wasted pollen and, in some cases, stigma clogging (Morales and Traveset 2008). Because of the intimate tie between pollinator behavior and reproduction, ethological isolation can be an important barrier to heterospecific mating (reviewed in Grant 1949; Kay and Sargent 2009; Van Der Niet et al. 2014). Despite numerous studies documenting pollinator mediated reproductive isolation in plants, we still lack a quantitative framework linking variation in pollinator behavior to variation in heterospecific pollen movement.

Pollinators impose strong selection on floral traits and it is thought that plants evolve to best utilize the most effective pollinator (reviewed in Fenster et al. 2004; Harder and Johnson 2009; Rosas-Guerrero et al. 2014; Schiestl and Johnson 2013). Pollinators can in turn cause reproductive isolation between closely related taxa. A number of case studies demonstrate the importance of pollinator behavior in causing reproductive isolation between sympatric species (*e.g.* Hopkins and Rausher 2012; Kay and Schemske 2003; Klahre et al. 2011; Ramsey et al. 2003), and reviews of reproductive isolation in plants more generally have found that pollinator mediated reproductive isolation has significantly contributed to speciation in plants (Baack et al. 2015; Lowry et al. 2008). Yet, there is still doubt as to the effectiveness of pollinators in preventing heterospecfic gene flow (Chittka et al. 1999; Waser 1998). Most plants are generalists appealing to a variety of pollinators, and most pollinators are generalists visiting a variety of plants (Jordano 1987; Ollerton 2016; Robertson 1928; Waser et al. 1996). How then can pollinators cause reproductive isolation? In order to better address this question we need a framework for evaluating how quantifiable properties of pollinator behavior lead to reproductive isolation in plants.

Two aspects of pollinator behavior – preference and constancy – influence reproductive isolation in plants. The relative importance of constancy and preference to plant reproductive isolation has been debated in the literature (Kay and Sargent 2009; Waser 1998) but a direct comparison has never been possible. There is currently no method for comparing the strength of preference to the strength of constancy and determining their respective contributions to heterospecific pollen movement.

Pollinator preference is the tendency of a pollinator to visit one species or variety of plant more than is expected based on that plant’s frequency in a population. Preference can be expressed in response to an assortment of traits, such as color, size, shape, and smell, which often act as signals for a reward such as nectar (Schiestl and Johnson 2013). Increased pollinator visitation due to preference is an important agent of selection implicated in the evolution and divergence of floral traits (*e.g.* Alexandersson and Johnson 2002; Benitez-Vieyra et al. 2006; Campbell et al. 1996; Mitchell et al. 1998). Preference can also cause reproductive isolation (*e.g.* Bradshaw and Schemske 2003; Fulton and Hodges 1999; Hoballah et al. 2007; Martin et al. 2008). In a community with two plant species, a pollinator that strongly prefers one species will have less opportunity to move pollen between species of plants than a pollinator that visits both species equally. For example, hummingbirds strongly prefer the red flowers of *Mimulus cardinalis* while bees prefer the pink flowers of *Mimulus lewisii* (Bradshaw and Schemske 2003). This pollinator specialization results in hummingbirds transferring pollen between *M. cardinalis* individuals, bees transfering pollen between *M. lewisii* individuals, and few heterospecific pollen transfers.

Pollinator constancy is the tendency of pollinators to move between the same species or variety of plant more than between different plants given what is expected based on the proportion of each plant visited (Waser 1986). Constancy describes the order of visits to plants rather than the number of visits to each type of plant. For example, in the Texas *Phlox* wildflower system, dark-red *P. drummondii* plants co-occur with the light-blue *P. cuspidata* plants and the two species share pollinators (Hopkins and Rausher 2012). Despite visiting both species of *Phlox,* the butterfly pollinators tend to move between plants of the same species more than plants of different species (Hopkins and Rausher 2012). In other words they go from a dark-red, to a dark-red, to a dark-red flower and then switch to go from a light-blue, to a light-blue, to a light-blue flower. In this way constancy causes reproductive isolation because pollen is transferred between conspecific individuals more than between heterospecific individuals. Foraging with constancy has been studied most in honeybees and bumblebees (Gegear and Laverty 2005; Hill et al. 1997; Kephart and Theiss 2004; Marques et al. 2007; Raine and Chittka 2005) but has been documented in Lepidoptera (Aldridge and Campbell 2007; Goulson and Cory 1993; Goulson et al. 1997; Hopkins and Rausher 2012; Kulkarni 1999; Lewis 1986) and hoverflies as well (Goulson and Wright 1998). Although, it is still unclear why pollinators forage with constancy, it has been hypothesized that it is an adaptive foraging strategy that optimizes efficiency of resource acquisition (Gegear and Thomson 2004; Goulson 1999).

Co-occurring plant species vary extensively in relative frequency over space and time. Intuitively, the proportion of heterospecific pollen transfers is correlated with the relative frequency of plant species. If a population is predominately made up of one species, just by chance there will be fewer opportunities for a pollinator to transfer pollen from the rare species to the common species than between the common species. But, we lack an understanding of how aspects of pollinator behavior, such as constancy and preference, affect this relationship between heterospecific pollen transfer and plant frequency.

Most plants are visited by multiple pollinators (Robertson 1928; Waser et al. 1996) that do not show the same behavioral responses to floral signals. How is reproductive isolation in plants affected if one pollinator has strong preference and another has weak preference? What if the frequency of these two pollinators varies across populations? A quantitative framework for understanding how multiple pollinators that express different behavior interact to cause reproductive isolation in plants is needed.

To address these gaps in our knowledge of pollinator mediated reproductive isolation I construct a mathematical model that describes how aspects of pollinator behavior cause heterospecific matings in plants. My specific goals are: 1.) To define a mathematical relationship between pollinator behavior (constancy and preference) and plant reproductive isolation, 2.) To determine how variation in plant frequency affects plant reproductive isolation and 3.) To determine how variation in pollinator frequency affects plant hybridization in a community with multiple pollinators. To demonstrate the relevance of this work to empirical investigations of plant-pollinator interactions, I apply this model to natural examples to predict proportion of heterospecific matings given preference and constancy measures taken from field trials with sympatric plants.

## MODEL

I present a deterministic model that describes how pollinator behavior leads to movement of pollen between conspecific and heterospecific plants. The notation used in my model is defined in Table 1. The goal of the model is to predict the proportion of heterospecific matings (*H*) between two co-occurring plant taxa, given pollinator behavior. This is a simplified model based on two parameters describing aspects of pollinator behavior – preference and constancy. The model investigates how the proportion of heterospecific matings varies with frequency of a pollinator and frequency of a plant. For simplicity I will start by describing a system with a single pollinator and two plants in which one plant is the focal species and the other is the heterospecific pollen donor. I will go on to describe a system with two pollinators and two plant species. Online Appendix A describes a generalized model that allows for unlimited plants and pollinators.

**Table 1.**
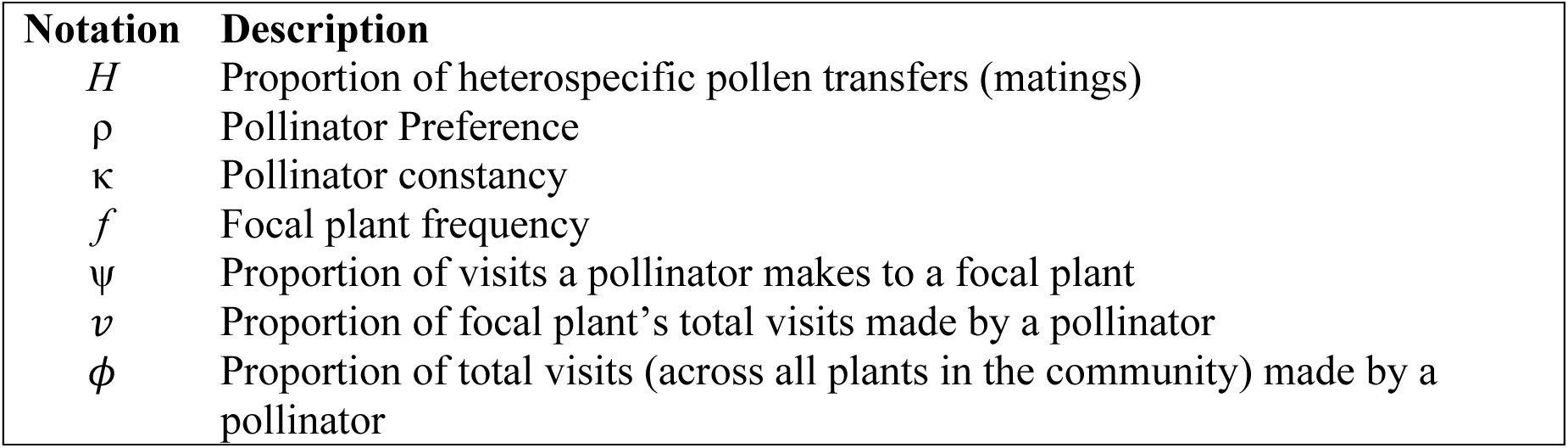
Summary of notation and definitions for model.

### One pollinator model

For a given pollinator the proportion of visits (ψ) to a focal plant can be described as a function of its preference (ρ) for the plant and the plant frequency (*f*).

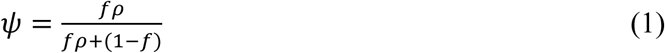

Equation 1 is modified from a foraging preference function in Greenwood and Elton (1979) as in Smithson and Macnair (1996) and can be adapted to include frequency dependent variation in preference (Online Appendix B). The proportion of visits to a plant species varies between 0 and 1 and increases as both frequency of the plant increases and preference for the plant increases. Preference varies from 0 to infinity and determines how much a pollinator over or under visits a plant given its frequency. If a pollinator shows no preference ρ = 1 and proportion of visits is equal to the frequency of the focal plant. If ρ > 1 a pollinator favors the focal species by visiting it ρ times more than is expected based on the plant frequency. In other words, if ρ = 2 a pollinator visits the focal species twice as much as the other plant given their frequencies. When ρ < 1 a pollinator disfavors the focal species and will visit it proportionally less. Preference for a given plant type is always relative to at least one other plant type in the population.

Pollinator constancy (*κ*) is a measure of the conspecific versus heterospecific transitions a pollinator makes given what is expected based on the frequency of visits to each plant. Constancy varies between −1 and 1, with negative constancy indicating more heterospecific transitions than expected and positive constancy indicating more conspecific transitions than expected. When observing pollinator movement, constancy can be calculated based on the observed proportion of heterospecific transfers (*H*) made by a pollinator to a plant and the observed proportion of visits to the focal species by the pollinator (ψ).

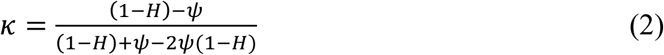

Equation 2 is modified from the constancy formula presented in Gegear and Thomson (2004). This equation can be solved for *H* in terms of constancy and proportion of visits. This yields

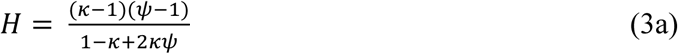

where the proportion of heterospecific pollen transfers is a function of preference and frequency of the focal plant. This can then be expanded to create an equation describing the proportion of heterospecific matings in terms of preference, constancy, and frequency of plants.

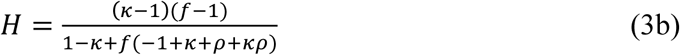

Equation 3b predicts the proportion of heterospecific pollen transfers a plant will experience given the preference and constancy of a pollinator.

#### The effect of preference

The proportion of heterospecific matings has an inverse relationship with focal plant frequency. The shape of this relationship depends on the strength of pollinator preference. With no constancy (*i.e.* expected number of conspecific transitions given frequency of visits),

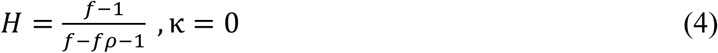

where the proportion of heterospecific matings is a function of pollinator preference and frequency of the focal plant. Figure 1a shows equation 4 evaluated for no preference, strong preference, weak preference, and preference against a focal plant.

The way in which plant frequency affects heterospecific matings depends on the strength of pollinator preference in the system. With no preference, *H* = 1 – *f*, such that the proportion of heterospecific matings is the inverse of the focal plant frequency. When a pollinator has preference for the focal species heterospecfic mating is a convex function of frequency. When a pollinator has preference against the focal species heterospecific mating becomes a concave function of frequency. The curvature of the function is determined by the strength of preference. When preference is strong (*ρ* = 10) and the focal plant is rare, small changes in plant frequency can result in significant declines in heterospecfic matings. When preference is weak (*ρ* = 2), the proportion of heterospecfic matings decreases less with an equivalent change in plant frequency. With strong preference against the focal species (*ρ* = 0.1), heterospecific matings remain frequent even at high focal plant frequencies and drop off precipitously as the focal plant becomes fixed in the population.

#### The effect of constancy

Similar to preference, a change in pollinator constancy will affect the way plant heterospecfic matings vary with plant frequency. In a system with no pollinator preference (ρ = 1),

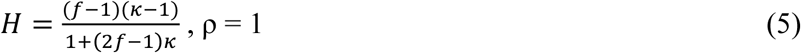

the proportion of heterospecific matings becomes a function of plant frequency and constancy. Figure 1b shows equation 5 evaluated for strong constancy, weak constancy, and strong negative constancy. Constancy causes heterospecific mating to be a convex function of plant frequency and negative constancy causes heterospecfic mating to be a concave function of plant frequency. The curvature of this function is dependent on the strength of constancy.

Equation 3b can be used to compare the relative importance of preference and constancy for plant reproductive isolation and how the two aspects of pollinator behavior interact. For example, Figure 1c shows how strong and weak preference and constancy interact. For these parameter values, a pollinator with strong preference and weak constancy is similar to a pollinator with strong constancy and weak preference.

**Figure 1:**
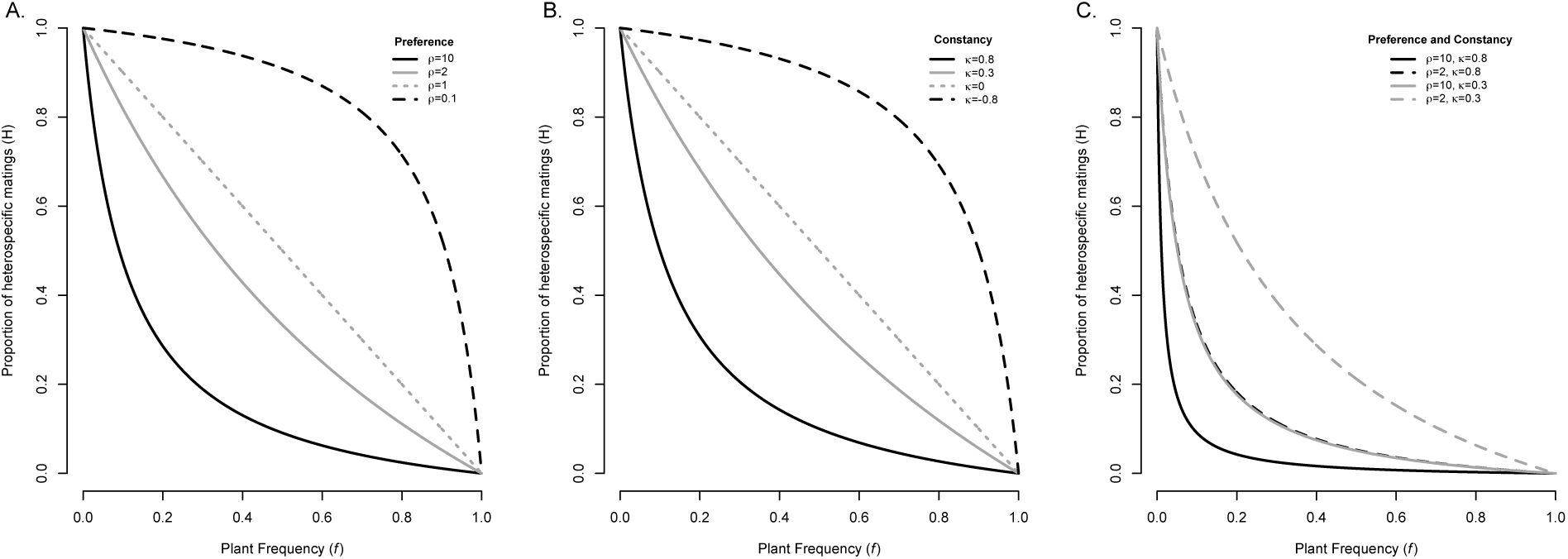
The relationship between plant frequency and proportion of heterospecific matings when pollinators have different preference and constancy. In A., examples of pollinators with no constancy (*κ* = 0) and strong preference for a plant (ρ =10, black line), weak preference for a plant (ρ=2, gray line), no preference for plant (ρ=1, dotted line), and strong preference against a plant (ρ=0.1, dashed line) show how the heterospecfic mating function is convex, linear, or concave depending on the strength and direction of preference. In B., examples of pollinators with no preference (ρ = 1) and strong constancy (κ = 0.8, black solid line), weak constancy (κ = 0.3, gray line), no constancy (κ = 0, dotted line), and negative constancy (κ =-0.8, dashed line) show how the heterospecific mating function depends on the strength of constancy. In C., examples of pollinators with strong preference and strong constancy (solid black line), weak preference and strong constancy (dashed black line); strong preference and weak constancy (solid gray line), and weak preference and weak constancy (dashed gray line) demonstrate how the affect of constancy and preference on heterospecific matings interact and can be compared.

#### Two pollinator model

Most plants receive visits from multiple pollinators that display different behavioral responses to floral traits. This observation motivates the need to understand how multiple pollinators with different strength preference and constancy combine to cause the total proportion of heterospecific matings a plant experiences. As mentioned above, I describe a model with two pollinators here and generalize this for any number of pollinators in Online Appendix A.

The total proportion of heterospecific pollen transferred to a plant is determined by the heterospecific pollen transferred by each pollinator proportional to the pollinator visitation rates. For a two pollinator system

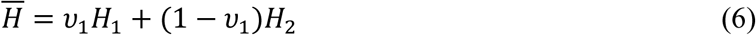

where 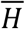 is the total proportion of heterospecific matings for a focal plant. In this equation, *H*_1_ and *H*_2_ are the proportion of heterospecific pollen transfers from pollinator 1 and pollinator 2 respectively, and *v*_1_ is the proportion of a plant’s total visits made by pollinator 1. I can define *v*_1_ as

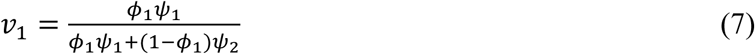

where *ϕ*_1_ is the frequency of visits by pollinator 1 across all plants. This term is related but distinct from pollinator frequency in a community because different types of pollinators may have different absolute numbers of visits. For the purpose of this model I am interested in frequency of visits across all plants. Additionally, this is distinct from *ψ*_1_ and *ψ*_2_, which are the proportion of visits pollinator 1 and 2 make to the focal plant. As above, *ψ*_1_ and *ψ*_2_ are functions of pollinator preference and plant frequency (equation 1). In this way the total proportion of heterospecific matings for a focal species can be predicted from pollinator behavior (preference and constancy), plant frequency, and pollinator visit frequency.

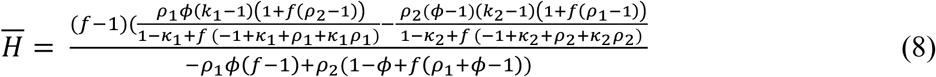

To explore the behavior of this equation I investigate variation in pollinator constancy and preference separately. For each aspect of behavior I show how the proportion of heterospecific matings varies across focal plant frequency when pollinator frequency is held at 50%, and across pollinator frequency when focal plant frequency is held at 50%.

#### Effect of preference across plant and pollinator frequency

First, I explore the way in which multiple pollinators that differ in preference affect the proportion of heterospecfic matings. If plant frequency is held at 50%, the proportion of heterospecfic matings is a nonlinear function of pollinator frequency (Figure 2a). This concave relationship reveals that the predicted proportion of heterospecific matings from two pollinators with different preference is not simply the average of the heterospecific matings by each pollinator, as this would predict a straight line. Instead, the pollinator with stronger preference for the focal plant disproportionally influences the total proportion of heterospecific matings.

**Figure 2:**
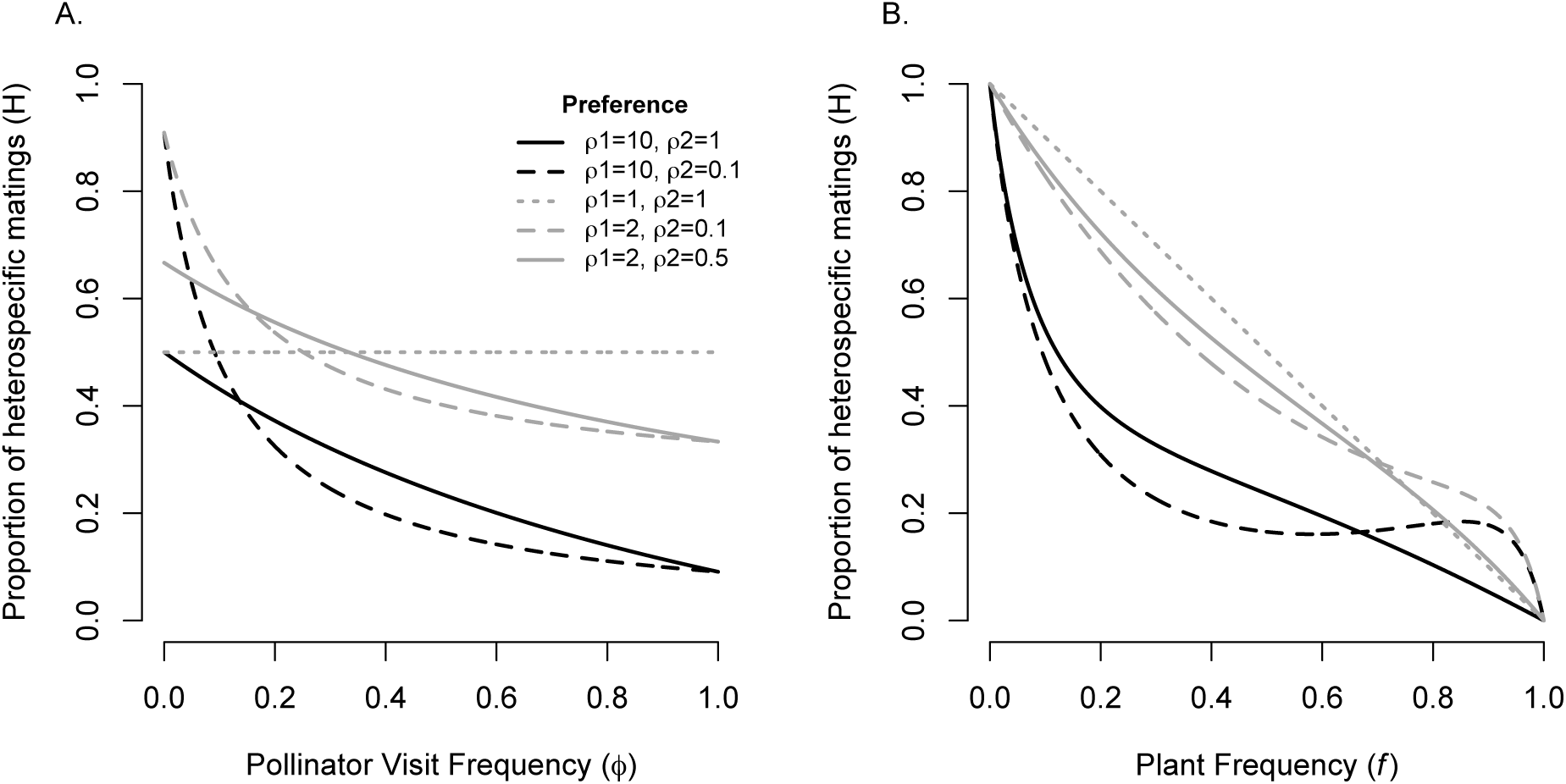
The relationship between plant frequency and proportion of heterospecific matings when two pollinators differ in preference across pollinator frequency (A.), or plant frequency (B.). Both plots show examples of pollinators with no constancy (κ = 0), but different strengths of preference. Black lines are scenarios with one pollinator having strong preference (ρ_1_ =10), and gray lines are when one pollinator has weak preference (ρ_1_=2) or no preference (ρ_1_= 1, dotted). Solid lines show when a second pollinator has no preference (ρ_2_ = 1, black) or weak preference against (ρ_2_ = 0.5, gray) and dashed lines show when a second pollinator has strong preference against the focal plant (ρ_2_=0.1). Note how dashed lines fall below solid lines across most pollinator frequencies.

The magnitude of this deviation from the linear average is proportional to the difference in preference between the two pollinators. This means that a plant receives fewer heterospecific matings when it has one pollinator that has strong preference against it, than if it has one pollinator with no preference at all (*i.e.* the black dashed line is below the black solid line) across most pollinator frequencies.

Second, I demonstrate the relationship between plant frequency and heterospecific matings when two pollinators are at equal frequency but differ in their preference (Figure 2b). I show a scenario for which one pollinator has strong preference and one pollinator has no preference for the focal plant (solid black line). This reveals a non-linear relationship between plant frequency and heterospecific matings such that at low focal-plant frequencies, small changes in plant frequency result in large decreases in heterospecfic matings. This rate of change in heterospecfic matings decreases as plant frequency increases. When one pollinator has strong preference for the focal plant and one pollinator that has strong preference against the focal plant (black dashed line), heterospecific mating becomes a non-monotonic function of plant frequency. Under some conditions heterospecific mating can actually increase as focal plant frequency increases. Although, across most plant frequencies (*i.e.* 0.3< *f* <0.9) the proportion of heterospecific matings changes very little. Of note, across many plant frequencies (*i.e* 0< *f* <0.67 in figure 2b) a focal plant has less heterospecfic matings when one pollinator has strong preference against the focal plant than if one pollinator has no preference at all. In Figure 2b this occurs when the dashed black and gray lines are below the solid black and gray lines.

#### Effect of constancy across plant and pollinator frequency

I explore the relationship between plant frequency and proportion of heterospecific matings when a plant has two pollinators that differ in the strength of constancy. If pollinators show no preference, (*ρ* = 1), then *v*_1_, the proportion of a plant’s total visits made by pollinator 1, equals the proportion of total visits pollinator 1 makes in the system (*ϕ*_1_). Therefore, the total proportion of heterospecific matings for a plant is the average of the proportion of heterospecific matings made by each butterfly weighted by the frequency of the butterfly in the system. Thus, in a two-pollinator system, the proportion of heterospecific matings varies linearly with pollinator frequency (Figure 3a). The slope of this line is determined by the difference in the proportion of heterospecific matings predicted by the two pollinators. Equation 6 can be rearranged to describe the linear relationship between total heterospecific matings and proportion of visits in the system by a pollinator:

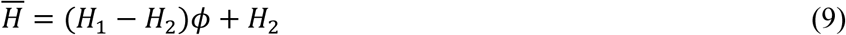

**Figure 3:**
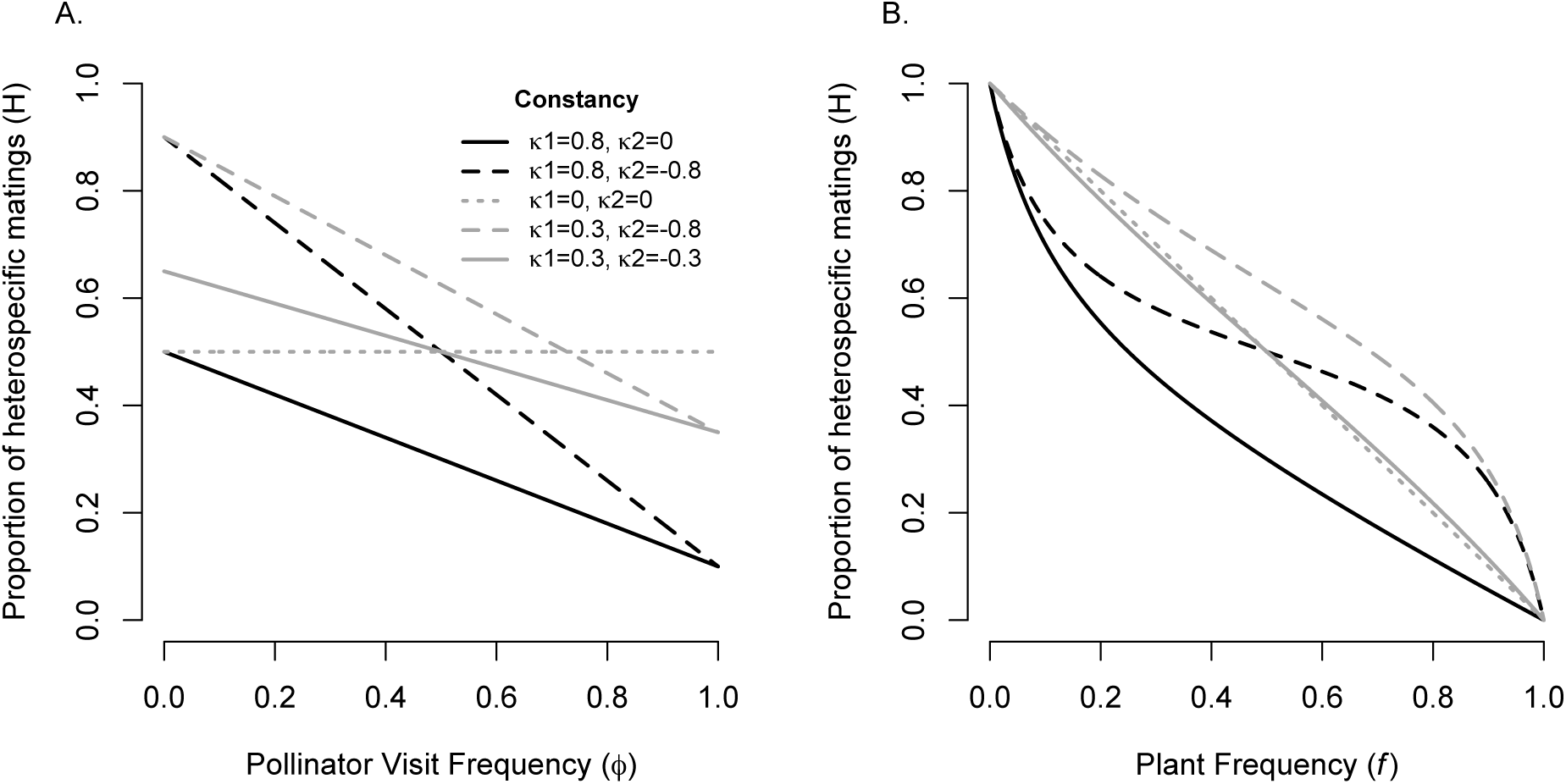
The relationship between plant frequency and proportion of heterospecific matings when two pollinators differ in constancy across pollinator frequency (A.), or plant frequency (B.). Both plots show examples of pollinators with no preference (ρ = 1), but different strengths of constancy. Black lines are scenarios with one pollinator having strong constancy (κ_1_ =0.8), and gray lines are when one pollinator has weak constancy (κ_1_=0.3) or no constancy (κ_1_= 0, dotted). Solid lines show when a second pollinator has no constancy (κ_2_ = 0, black) or weak negative constancy (κ_2_ = −0.3, gray) and dashed line shows when a second pollinator has strong negative constancy (κ_2_=-0.8).

In figure 3b, I show the relationship between pollinator frequency and proportion of heterospecific matings when two pollinators differ in the strength of constancy. In a two-pollinator system, the total proportion of heterospecific matings is the average of the proportions predicted by each pollinator independently. Recall that the relationship between heterospecfic matings and plant frequency is non-linear in a one-pollinator system therefore it is also nonlinear in a two-pollinator system. Note, unlike with preference variation, a plant always has fewer heterospecific matings when one of its pollinators has no constancy than when a pollinator has negative constancy.

### Model applications

Below I apply this mathematical model to empirical data of pollinator behavior. Details of how to identify and calculate the necessary parameters from field observations are described in Online Appendix C.

#### Pollinator preference and constancy for Phlox

Previous work shows that pollinator behavior is a significant reproductive isolating barrier between sympatric *P. drummondii* and *P. cuspidata* (Hopkins et al. 2014). Flower color divergence in *P. drummondii* is due to reinforcement. A change in flower color evolved in sympatric populations because the derived color decreased heterospecific matings between these two *Phlox* species (Hopkins et al. 2014). Specifically, when both *Phlox* had the ancestral light-blue flower color the pollinator *Battusphilinor* (the Pipevine Swallowtail) moved pollen extensively between the two species, but when *P. drummondii* had dark-red flower color, the butterfly made significantly fewer transitions between species and more transitions within species (Hopkins et al. 2014). These pollinator observations were done on arrays of plants at fixed frequency (60% *P. cuspidata,* 20% dark-red *P. drummondii*, and 20% light-blue *P. drummondii*). The observations indicate that the change in flower color can decrease heterospecific matings by 50%, but how do the proportion of heterospecific matings for light-blue and *P.drummondii*). The observations indicatethat the change in flower color candecrease heterospecific matings by 50%,but how do the proportion of heterospecific matings for light-blue and dark-red plants vary across different frequencies of *Phlox* as you might expect to find in the wild? I use equation 3b to evaluate the expected proportion of heterospecific matings for light-blue and dark-red *P. drummondii* across plant frequencies given *B. philenor* behavior (Figure 4). Based on the field observations of *B. philenor,* I used equation 1 and equation 2 to calculate preference and constancy, respectively. Across all *P. drummondii* frequencies, dark-red flowers are expected to have a lower proportion of heterospecific matings than light-blue flowers. As noted previously, this difference in predicted heterospecific matings is due to variation in pollinator constancy based on flower color. The expected difference between the two colors decreases at both high and low frequencies. Since the strength of reinforcing selection is proportional to the difference in heterospecific matings between the two flower colors, these results suggest that the strength of reinforcing selection is dependent on the frequency of the two species of plants in the population.

**Figure 4:**
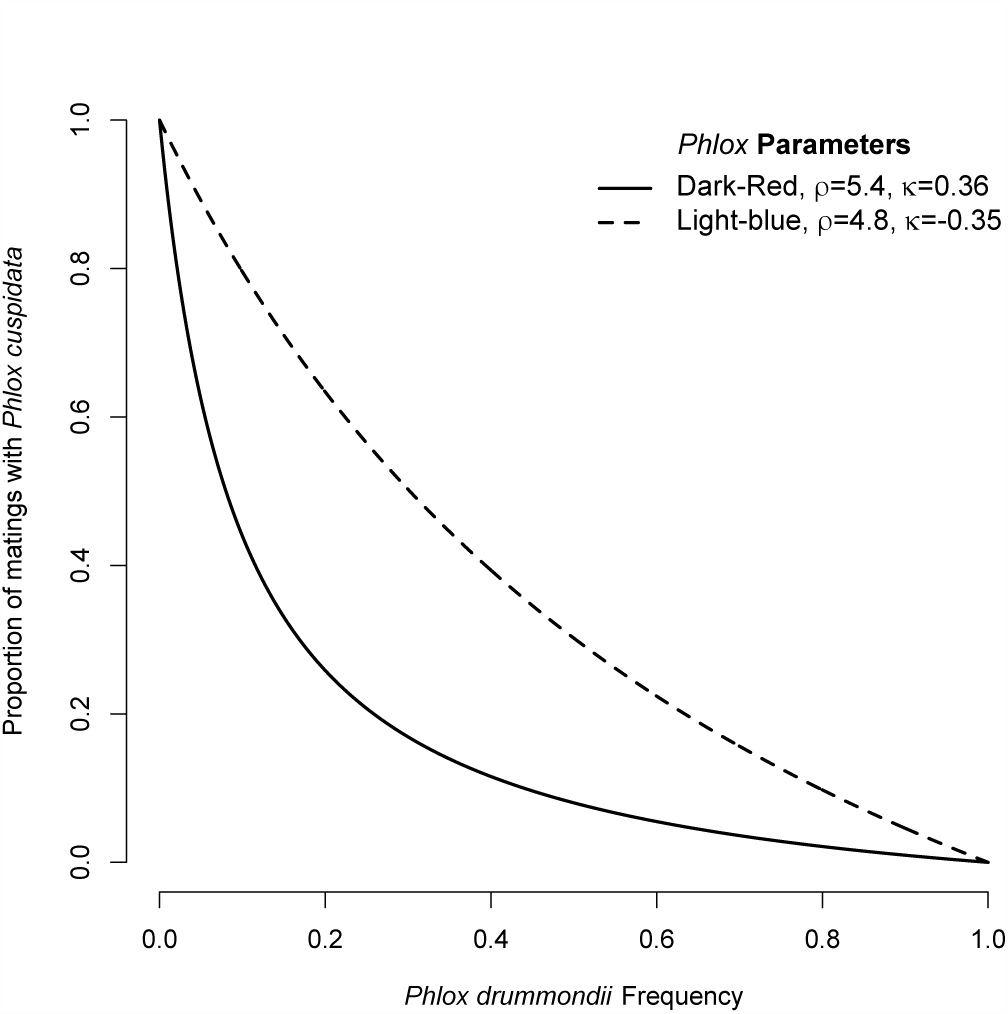
Predicted proportion of heterospecific matings between *Phlox drummondii* and *Phlox cuspidata* across *P.drummondii* frequency. Preference and constancy were estimated from field observations described by Hopkins et al(2014). Equation 3b was used to evaluate proportion of heterospecific matings

#### Two pollinators’ preferences for Mimulus

A classically studied species pair for which pollinator behavior is known to play an important role in reproductive isolation is *Mimulus cardinalis* and *Mimulus lewisii* (Ramsey et al. 2003). These two species differ in a number of floral traits that influence pollinator preference such that hummingbirds strongly prefer *M. cardinalis* and bees strongly prefer *M. lewisii* (Bradshaw and Schemske 2003; Byers et al. 2014; Ramsey et al. 2003; Schemske and Bradshaw 1999). Based on the pollinator observation data reported by Ramsey et al. (2003) on natural populations of the two *Mimulus* species, I quantified pollinator preference from the number of visits to each species by each pollinator and the frequency of each plant species. These two species grow across an elevation cline with *M. cardinalis* tending to grow more at lower elevations and *M. lewisii* found at higher elevation (Ramsey et al. 2003). Across this cline there is presumably variation in the relative frequency of the two species. I evaluate how heterospecifc matings between these two species is predicted to change across plant frequency given the pollinator preference observed in the field. Figure 4 shows equation 8 evaluated across frequencies of *M. cardinalis* for both species of *Mimulus.* As expected given the strong preference observed, reproductive isolation is nearly complete for both *M. cardinalis* (solid line) and *M. lewisii* (dashed line). The simplest form of my model, assuming no frequency dependent change in preference, predicts a higher proportion of heterospecific matings at the edges of the cline when one of the two species is rare. Intuitively, one might expect the highest rate of hybridization to be when both species are coexisting at equal frequency. In fact this is exactly the opposite of what my model predicts. At high frequencies, *M. cardinalis* is predicted to suffer from a relatively high proportion of heterospecific matings. Although bees prefer *M. lewisii,* they have been observed visiting *M. cardinalis* (Schemske and Bradshaw 1999). The model predicts that when *M. cardinalis* is frequent, the preferred visits to *M. lewisii* will be interspersed by visits to *M. cardinalis* resulting in heterospecific transfer of pollen.

**Figure 5:**
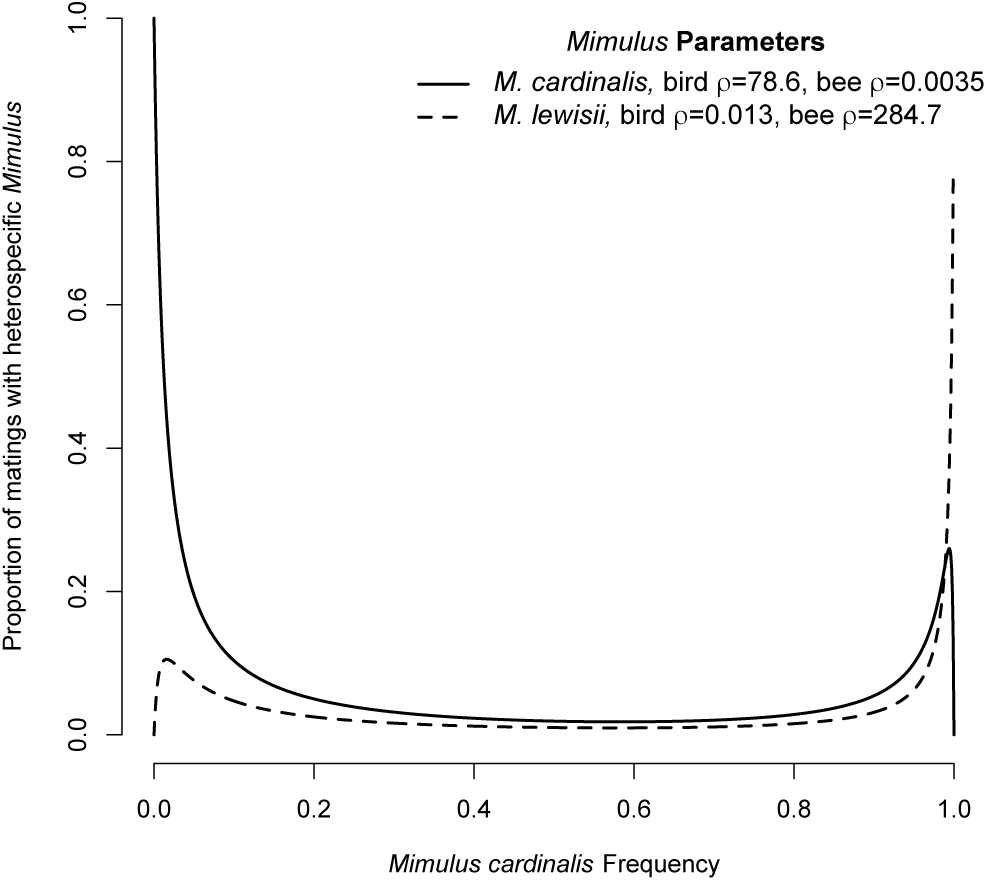
Predicted proportion of heterospecific matings between *M. cardinalis* and *M. lewisii.* Preference for both hummingbirds (bird) and bees were estimated from field observations described by Ramsey et al (2003). Equation 8 was used to evaluate proportion of heterospecific matings.

During this experiment humming birds visited 138 plants but none of them were *M. lewisii* indicating very strong preference. Unfortunately, using equation 1, preference is infinite when no visits to the second species are observed. To overcome this problem I estimated the proportion of visits to *M. cardinalis* to be exceedingly high (ψ = 0.996). Other studies have documented hummingbirds visiting *M. lewisii* indicating this estimate is not unreasonable (Schemske and Bradshaw 1999). Of note, the shapes of the curves in Figure 4 do not qualitatively change when preference is increased ten-fold, indicating that if preference is very strong the model is not sensitive to error in preference measurements.

## DISCUSSION

My model predicts how reproductive isolation caused by pollinator behavior varies across plant-pollinator communities. Specifically, I developed a quantitative framework for understanding how the frequency of hybridizing plants and the frequency of pollinators affects pollinator mediated reproductive isolation in plants. This research focuses on two important aspects of pollinator behavior – preference and constancy – that describe which plants a pollinator visits and the order of plant visitation. Both aspects of behavior have been shown empirically to contribute to plant speciation (Fenster et al. 2004; Kay and Sargent 2009), but this is the first model that describes how variation in behavior leads to variation in the proportion of heterospecific matings.

### Model implications

My research is motivated by the observation that plant-pollinator communities vary over geographic space and yet we lack a clear understanding of the implications of this variation for plant reproductive isolation. As discussed below, other authors have addressed this problem in a variety of ways. I took an analytical approach to derive the simplest model based on quantifiable aspects of pollinator behavior, frequency of plant species, and frequency of pollinators in the system to predict the proportion of heterospecific matings a plant will experience in a particular community.

My model reveals some intuitive results – for example, reproductive isolation increases as pollinator preference and constancy increase – as well as some non-intuitive results. I found that the proportion of heterospecific matings (the inverse of reproductive isolation) decreases as focal plant frequency increases in a community. The relationship between heterospecific matings and plant frequency is non-linear and the shape of this curve is determined by the strength of pollinator preference and constancy. I found that the stronger the behavior, the steeper the curve, such that strong preference or constancy leads to low heterospecific matings even at low focal plant frequencies.

The relationship between plant frequency and proportion of heterospecific matings is more complex in a multi-pollinator community. It has long been assumed, and observed, that if two closely related plant species attract different pollinators that express opposing preferences, than this specialization will result in reproductive isolation between the plants (Fenster et al. 2004; Kay and Sargent 2009). The advantage of pollinator specialization has been an emergent property in other theoretical models describing plant-pollinator communities (Sargent and Otto 2006), but for the first time, I reveal how pollinator behavior causes this advantage. When multiple pollinators have different preferences, their relative foraging efforts in a plant community influence the predicted total proportion of heterospecific matings a focal plant experiences. Yet, the pollinator with stronger preference has a disproportionately large influence on the total proportion of heterospecific matings regardless of its frequency. In fact, the more different two pollinators are in the strength of their preferences, the more total heterospecific matings reflects the behavior of the pollinator with a stronger preference. This means that across most plant and pollinator frequencies, a focal plant will experience a lower total proportion of heterospecific matings if one pollinator prefers the focal plant and a second pollinator has preference against the focal plant than if the second pollinator has no preference at all. In other words, my model demonstrates that pollinator specialization is favored because pollinators with different preferences interact non-additively to give the total proportion of heterospecific matings a plant will experience.

Unlike with preference, constancy from two different pollinators does combine additively to predict the total heterospecific matings. If two pollinators differ in the strength of their constancy, then the total proportion of heterospecific matings predicted by these two pollinators is the average heterospecific matings predicted by each pollinator weighted by the relative frequency of foraging visits made by each pollinator in the community.

### Model applications

The results from my model provide expectations for how pollinator mediated reproductive isolation should vary across populations that differ in plant and pollinator composition. A strength of my model is that empirical observations of pollinator behavior in natural or artificial plant communities can be used to predict pollinator mediated reproductive isolation across a landscape. Specifically, pollinator preference can be calculated from observations of proportion of visits a pollinator makes to multiple floral types and the proportion of those floral types available. Pollinator constancy can be calculated from observations of the pattern of movements pollinators make and the proportion of visits a pollinator makes to each floral type. From these simple observations this model can predict reproductive isolation for any pollinator frequency or plant frequency that occurs in nature. In Online Appendix C, I describe specific examples of how this model can be applied using observations in *Phlox* and the *Mimulus.* Because of the ease of applying this model to empirical data, my model can provide novel insights into natural plant-pollinator communities. For example, it can explain why and how hybridization and gene flow vary across natural plant hybrid zones. This model can also make predictions about how hybridization will increase or decrease as climate change shifts plant and pollinator ranges, phenology, and population sizes (Memmott et al. 2007).

The predictions from my model are based on simplifying assumptions including: 1.) Pollinator preference and constancy are constant, 2.) The relevant spatial scale of plant populations and pollinator foraging populations align, 3.) There are no spatially dependent aspects of pollinator foraging within the relevant scale of inquiry, 4.) Effectiveness of pollen transfer is similar across pollinators, 5.) Plants are not pollinator limited such that heterospecific matings are replaced with conspecific matings. It is likely that natural plant-pollinator communities are more complicated than described by my model and violate some of these assumptions. For example, some pollinators show frequency dependent preference (*e.g.* (Cresswell and Galen 1991; Gigord et al. 2001; Smithson and Macnair 1996; Smithson and MacNair 1997) and, as seen in Online Appendix A, this changes the predicted proportion of heterospecific matings across populations. In other systems pollinators may vary their behavior depending on the behavior and frequency of other pollinators in the community (*e.g.* (Brosi and Briggs 2013; Fontaine et al. 2008; Inouye 1978)). The simplified model presented here can act as a null against which variation in natural systems can be tested. In other words, if natural variation in heterospecific matings does not conform to the expectations of the model, we now have a powerful framework with which to test alternative hypotheses about what other factors affect the proportion of heterospecific matings.

My model focuses specifically on the proportion of heterospecific matings and does not consider total fitness of a plant. In other words, the model assumes that there are always enough pollinators to pollinate flowers, and it is just a question of whether the delivered pollen is conspecific or heterospecific. In many systems this assumption is violated (Ashman et al. 2004; Burd 1994; Larson and Barrett 2000). I acknowledge that in a pollinator limited system the proportion of heterospecific matings a plant experiences is not the only, or maybe even the most important, source of selection. But, it is beyond the scope of this project to investigate how frequency of plants and pollinators might affect total plant fitness if there is pollinator limitation and heterospecific matings. Extending the model to incorporate such scenarios is an important future direction.

Previous theoretical research has considered aspects of plant-pollinator community context to understand pollinator specialization, pollinator network structure (Bascompte et al.2006), and how perturbations disrupt pollination (Kaiser-Bunbury et al. 2010; Memmott et al. 2004; Ramos-Jiliberto et al. 2012). Models such as these provide important insights into how and why plant-pollinator communities might be structured as they are and how their current structure can maintain biodiversity and stability. Some of these models have even incorporated explicit aspects of pollinator behavior, such as adaptive foraging to reveal further insights (Valdovinos et al. 2016; Valdovinos et al. 2013).

### Conclusions

My model is unique in its reductionist view. My goal is to deconstruct the quantifiable aspects of pollinator behavior to understand how variation in behavior contributes to reproductive isolation across different plant communities. The model can generate predictions about how temporal and spatial fluctuations in pollinator composition and plant composition influences pollinator mediated reproductive isolation. This can be useful for empiricists to better understand how the pollinator behavior they measure in the field leads to reproductive isolation across changing communities. It is also useful for researchers studying plant speciation to understand how components of reproductive isolation may vary across different populations. Finally, this model is worthwhile at a theoretical level in that it brings together commonly used, yet seemingly unrelated, equations for pollinator behavior and reproductive isolation to analytically describe ethological isolation in plants.

## Online Appendix A: Generalized model for multiple pollinators and multiple plants

Equation 6 for total heterospecific matings generalized for *n* pollinators is

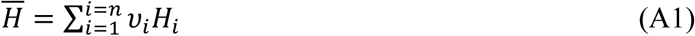

where *H_i_* is the proportion of heterospecific matings predicted for pollinator *i* and *v_i_* is the proportion of total plants visits made by pollinator *i*. Note that 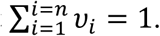

The equation describing the heterospecific matings for each pollinator is

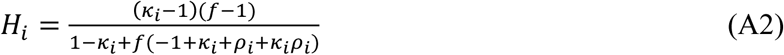

where *k_i_* and *ρ_i_* are constancy and preference of pollinator *i* for the focal plant.

Equation 7 for the total proportion of visits to a plant made by any pollinator *i* generalized for *n* pollinators is

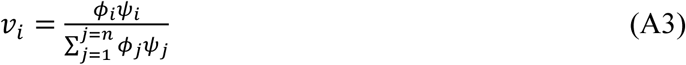

where *ϕ_i_* is the frequency of visits by pollinator *i* across all plants, such that the sum of all *ϕ_i_* across pollinators 1 through *n* equals 1. For each pollinator, *ψ_j_* is the proportion of visits it makes to just the focal plant.

The total proportion of heterospecific matings for a give focal plant can be predicted from the preference and constancy of each pollinators in a community with the following equation

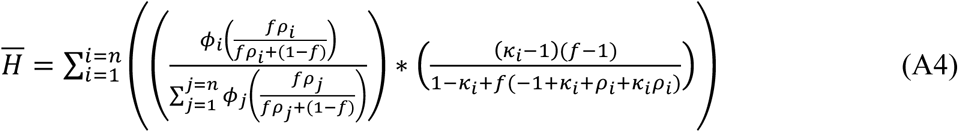

## Online Appendix B: Model of frequency dependent pollinator behavior

Frequency dependent changes in pollinator preference affect the predicted proportion of visits by a pollinator to the focal plant (ψ). The below equation describing proportion of visits by a pollinator includes a frequency dependent parameter (*b*), which is the coefficient of frequency-dependence as in Smithson and Macnair (1996):

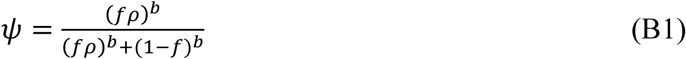

When *b* =1 there is no frequency dependent change in preference, when *b*>1 preference is stronger for more frequent plants representing positive frequency-dependence, and when *b*<1 preference is less for frequent plants representing negative frequency-dependence.

The proportion of heterospecific matings a focal plant is predicted to experience for a single pollinator *i* is

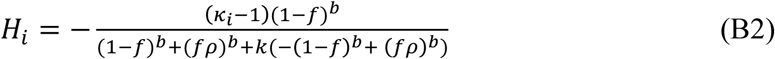

The total heterospecific matings for a focal plant given *n* pollinators and frequency dependent preference is

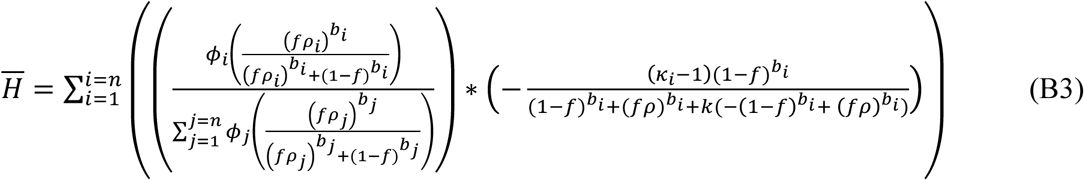

## Online Appendix C: Using the model with empirical data

### Phlox example with variation in constancy

Pollinator observations of *Battus philenor* were performed on arrays of *Phlox* plants (Hopkins et al. 2014). These arrays contained 10 light-blue *P. drummondii* plants, 10 dark-red *P. drummondii* plants and 31 *P. cuspidata* plants. Both the number of flower types visited by the pollinators and the number of transitions between each flower type was observed. I calculated proportion of heterospecific matings separately for light-blue and dark-red *P. drummondii* and therefore considered only the light-blue and *P. cuspidata* observations when calculating light-blue *P. drummondii* proportion of heterospecific matings and only dark-red and *P. cuspidata* observations when calculating dark-red *P. drummondii* heterospecific matings. Below I first describe how to use the observed data to calculate the model parameters in order to predict proportion of heterospecific matings for light-blue *P. drummondii* across all plant frequencies.

- The frequency of focal plant (light-blue *P. drummondii*) is given.

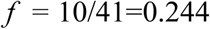
- The total visits to light-blue *P. drummondii* (300) and *P. cuspidata* (172) are used to calculate ψ (the proportion of visits to focal plant).

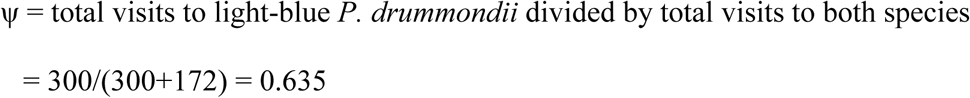
- Preference (ρ) is calculated from proportion of visits to focal plant (ρ) and frequency of focal plant (*f*).This is a rearrangement of equation 1.

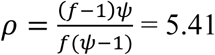
- Proportion of heterospecific matings (H) observed is calculated from observed pollinator transitions to light-blue *P. drummondii* from light-blue *P. drummondii* (73) and to light-blue *P. drummondii* from *P. cuspidata* (87).

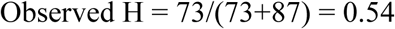
- Constancy (*k*) is calculated using equation 2.

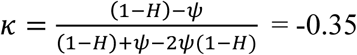

With preference (ρ) and constancy (*κ*), the model can be used to predict heterospecific mating for any frequency of light-blue *P. drummondii* and *P. cuspidata.*

Now I describe how to use the observed data to calculate the model parameters to predict proportion of heterospecific matings for dark-red *P. drummondii* across all plant frequencies.

- The frequency of focal plant (dark-red *P. drummondii*) is given.

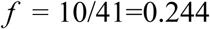
- The total visits to dark-red *P. drummondii* (266) and *P. cuspidata* (172) are used to calculate ψ (the proportion of visits to focal plant).

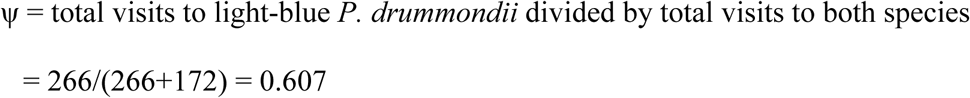
- Preference (ρ) is calculated from proportion of visits to focal plant (ψ) and frequency of focal plant (*f*).This is a rearrangement of equation 1.

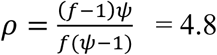
- Proportion of heterospecific matings (H) observed is calculated from transitions to light-blue *P. drummondii* from light-blue *P. drummondii* (95) and to light-blue *P. drummondii* from *P. cuspidata* (29).

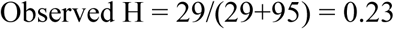
- Constancy (*k*) is calculated using equation 2.

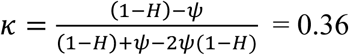

With preference (ρ) and constancy (*κ*), the model can be used to predict heterospecific mating for any frequency of dark-red *P. drummondii* and *P. cuspidata*.

### Mimulus example with preference varying across two pollinators

Pollinators were observed visiting mixed populations of *Mimulus cardinalis* and *M. lewisii* (Ramsey et al. 2003). The identity of the pollinator (bee *vs.* hummingbird), the plant visited, and the number of open flowers of each species were recorded. Two populations were observed but most data came from one population so the proportion of open flowers from that population will be used to calculated preference. The following data were observed:

- On average there were 12.8 open *M. lewisii* flowers and 40.6 open *M. cardinalis* flowers, which can be used to calculate the frequency of the focal plant (*f*).

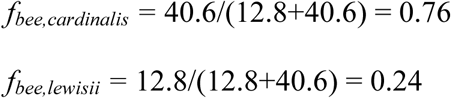
- Bees visited 259 *M. lewisii* flowers and 3 *M. cardinalis* flowers, which can be used to calculated proportion of visits to focal plant (ψ) from bee

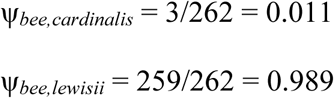
- Bee preference (ρ) for the two flowers can be calculated from the frequency and proportion of visits

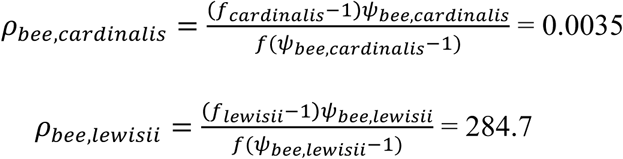
- Hummingbirds visited 138 *M. cardinalis* flowers and no *M. lewisii* flowers. In order to calculate proportion of visits and preference I used 0.5 as the number of *M. lewisii* flower visits.

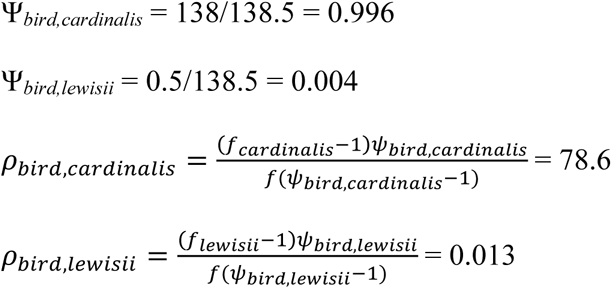
- In total, hummingbirds visited 138 flowers and bees visited 262 flowers. From this I can calculate ϕ, the proportion of visits to all plants made by each pollinator

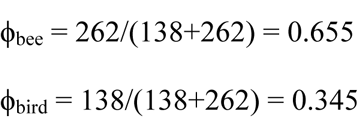

With preference (ρ) for each pollinator and the frequency of visits by each pollinator in the system (ϕ), the proportion of heterospecific matings for any *M. cardinalis* frequency can be predicted using equation 8.

